# Linked annotations: a middle ground for manual curation of biomedical databases and text corpora

**DOI:** 10.1101/014274

**Authors:** Tatyana Goldberg, Shrikant Vinchurkar, Juan Miguel Cejuela, Lars Juhl Jensen, Burkhard Rost

**Affiliations:** Bioinformatics & Computational Biology, Department of Informatics, Technical University of Munich (TUM), 85748 Garching, Germany; TUM Graduate School, Center of Doctoral Studies in Informatics and its Applications (CeDoSIA), 85748 Garching, Germany; Novo Nordisk Foundation Center for Protein Research, Faculty of Health and Medical Sciences, University of Copenhagen, 2200 Copenhagen N, Denmark

## Abstract

Annotators of text corpora and biomedical databases carry out the same labor-intensive task to manually extract structured data from unstructured text. Tasks are needlessly repeated because text corpora are widely scattered. We envision that a *linked annotation resource* unifying many corpora could be a game changer. Such an open forum will help focus on novel annotations and on optimally benefiting from the energy of many experts. As proof-of-concept, we annotated protein subcellular localization in 100 abstracts cited by UniProtKB. The detailed comparison between our new corpus and the original UniProtKB annotations revealed sustained novel annotations for 42% of the entries (proteins). In a unified linked annotation resource these could immediately extend the utility of text corpora beyond the text-mining community. Our example motivates the central idea that linked annotations from text corpora can complement database annotations.

## Background

The natural language processing (NLP) and biomedical research communities have in common that they invest great effort into making high-quality manual annotation of biomedical literature. The focus and the annotation strategies of the two communities have, however, differed so much that collaborations remained stunningly limited. Most text corpora contain detailed markup of only a few types of entities and relationships in a limited number of abstracts or articles [Neves, 2014] (with exceptions such as the CRAFT corpus [Verspoor et al., 2012]). In contrast, manually curated databases such as Swiss-Prot/UniProtKB [UniProt Consortium, 2014] aim at annotating each entity with a wide range of information extracted from literature, but with less focus on the text structure.

We envision *linked annotations* as a possible middle ground for the two important strategies to curate literature that could synergistically link the efforts of two distinct communities. By connecting the annotations of different types of entities and relationships annotated in existing and future corpora, a *linked annotation resource* could be constructed, which would have much greater coverage and diversity of annotations than any existing text corpus. Such a corpus would be valuable to NLP researchers and database curators alike.

Here, we present a case study on protein subcellular localization to demonstrate that the corpus annotation strategy can improve database annotation. The localization of a protein is one aspect of protein function and therefore constitutes one of the three hierarchies to capture protein function employed by the Gene Ontology (GO) [Ashburner et al., 2000].

## The LocText corpus

We assembled a corpus of 100 PubMed abstracts referenced by UniProtKB. We focused on three *model* organisms: *Homo sapiens* (50 entries), *Saccharomyces cerevisiae* (baker’s yeast with 25 entries), and *Arabidopsis thaliana* as a plant (25 entries). We used 46 of the 100 abstracts to develop our annotation guidelines that are available at https://www.tagtog.net/-corpora/loctext.

Two of us (TG & SV) then annotated the remaining 54 abstracts. The two annotations agreed at F_1_=94% for entities and at F_1_=80% for relationships. We normalized protein names to UniProtKB and localizations to GO identifiers. The resulting corpus contains 306 annotated relationships in 201 different UniProtKB proteins with 48 GO distinct localization terms. All annotations were made within the framework of the *tagtog* system (Figure 1) [Cejuela et al., 2014; http://tagtog.net] and Reflect was used to aid protein name normalization [Pafilis et al, 2009; http://reflect.ws]. The corpus is available for download at https://www.tagtog.net/-corpora/loctext under the Creative Commons Attribution 4.0 (CC-BY 4.0) license.

**Figure 1.**
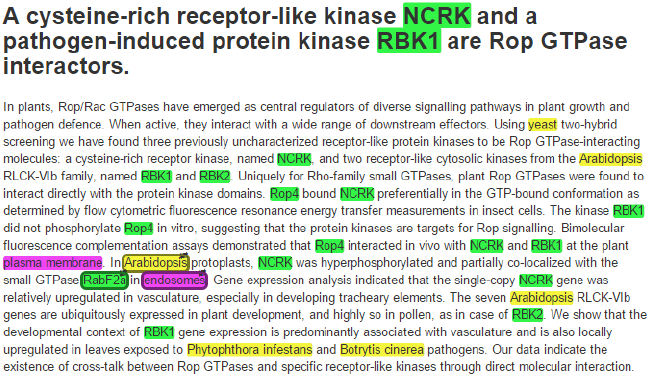
Curation of protein subcellular localization. The simplified *tagtog* web interface shown assisted in the manual annotation of the corpus (abstract of [Molendijk et al., 2008]). Colours highlight names of organisms (yellow), genes/proteins (green), and localization terms (magenta). Linking the *Arabidopsis* protein RabF2a (UniProtKB ID: RAF2A_ARATH) to endosomes adds a novel annotation to UniProtKB.

## Corpus provides novel annotations

Linked annotations from text corpora can complement database annotations only if manual corpus annotations identify relationships not captured by existing databases. Therefore, all our annotations were done from scratch without using database annotations. Comparing our “from scratch” annotations with those from UniProtKB revealed important novelty added by our text corpus.

We found novel or more detailed localization annotations with respect to UniProtKB for 84 of 201 (42%) proteins in 34 abstracts (Table 1); for example, *Arabidopsis* RabF2a (UniProtKB entry RAF2A_ARATH) is localized to endosomes (Figure 1). We found that for over half of these proteins with additional annotations (47/84=56%) UniProtKB did not cite the abstracts. This is likely explained by the way proteins are annotated, one protein at a time: if a curator works on one protein and an abstract mentions also the localization of another, which is not the focus of curator, the localization of the latter might not be annotated.

**Table 1.**
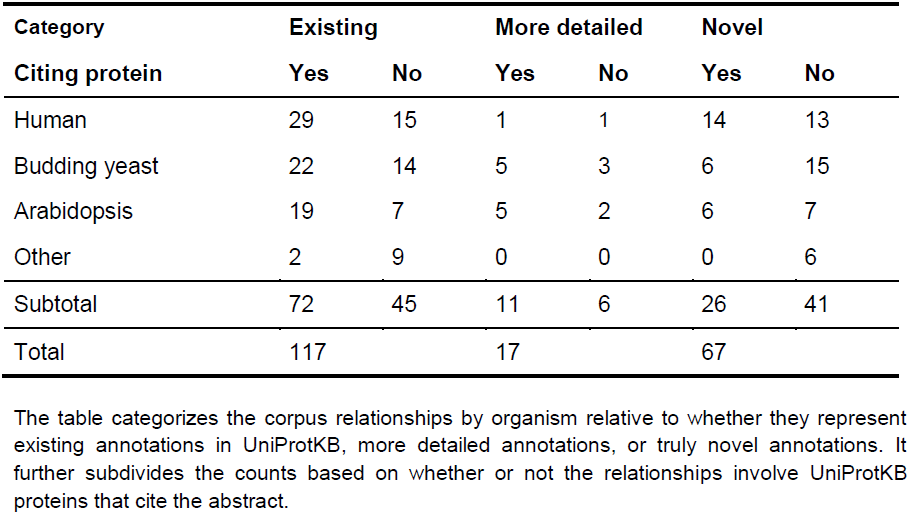
Localization annotations in our corpus and in UniProtKB.

## Perspectives

Our case study clearly showed that corpora containing manual annotations of the sub-cellular localization of proteins are able to contribute novel information to curated databases such as UniProtKB. Notably, this is even true in the worst-case example when limiting annotations only to abstracts of articles that have already been utilized by the database curators. We expect our findings to generalize to most types of protein annotation, including disease associations and tissue expression.

Today databases avoid the trouble of integrating these annotations, because most text corpora are too limited in size and scope. Having the corpus developers combine their annotations into a single, unified linked annotation resource could thus be an important step towards integration of corpus annotations into databases, thus making them to richer data collection systems. Even before integration with databases happens, it will be possible for researchers to use semantic web technologies to combine the information in the linked annotation resource with that in existing databases, since UniProtKB and many other databases are already Resource Description Framework (RDF) compliant.

We envision a linked annotation resource to continuously grow, supported by annotation tools making it easy for corpus developers to link future annotations; for example, through a standard JSON format. Not all linked annotations need to be made manually, though. Including also results from automatic text mining pipelines would help address the challenge of the prohibitively high costs of large-scale manual annotation [Baumgartner et al., 2007]. Associations extracted from both open and non-open access journals can be linked, as redistribution of extracted facts is not prohibited by most publishers’ licenses.

## Acknowledgments

Funding: Alexander von Humboldt Foundation through German Federal Ministry for Education and Research, Ernst Ludwig Ehrlich Studienwerk, and the Novo Nordisk Foundation Center for Protein Research (NNF14CC0001).

